# Dynamic changes in β-cell electrical activity and [Ca^2+^] regulates NFATc3 activation and downstream gene transcription

**DOI:** 10.1101/2020.06.28.176768

**Authors:** Jose G. Miranda, Wolfgang E Schleicher, David G. Ramirez, Samantha P Landgrave, Richard KP Benninger

## Abstract

Diabetes results from insufficient insulin secretion as a result of dysfunction to β-cells within the islet of Langerhans. Elevated glucose causes β-cell membrane depolarization and action potential generation, voltage gated Ca^2+^ channel activation and oscillations in free-Ca^2+^ activity ([Ca^2+^]), triggering insulin release. Nuclear Factor of Activated T-cell (NFAT) is a transcription factor that is regulated by increases in [Ca^2+^] and calceineurin (CaN) activation. NFAT regulation links cell activity with gene transcription in many systems, and within the β-cell regulates proliferation and insulin granule biogenesis. However the link between the regulation of β-cell electrical activity and oscillatory [Ca^2+^], with NFAT activation and downstream transcription is poorly understood. In this study we tested whether dynamic changes to β-cell electrical activity and [Ca^2+^] regulates NFAT activation and downstream transcription. In cell lines, mouse islets and human islets, including those from donors with type2 diabetes, we applied both agonists/antagonists of ion channels together with optogenetics to modulate β-cell electrical activity. Both glucose-induced membrane depolarization and optogenetic-stimulation triggered NFAT activation, and increased transcription of NFAT targets and intermediate early genes (IEGs). Importantly only conditions in which slow sustained [Ca^2+^] oscillations were generated led to NFAT activation and downstream transcription. In contrast in human islets from donors with type2 diabetes NFAT activation by glucose was diminished, but rescued upon pharmacological stimulation of electrical activity. Thus, we gain insight into the specific patterns of electrical activity that regulate NFAT activation and gene transcription and how this is disrupted in diabetes.

## Introduction

The β-cell plays a crucial role in maintaining glucose homeostasis by secreting insulin, which promotes glucose uptake and maintains glucose homeostasis. Diabetes, a disease afflicting >400M people worldwide generally results from dysfunction or death to β-cells and insufficient release of insulin. Following elevations in blood glucose levels, the β-cell undergoes a series of steps that includes metabolism of glucose to elevate ATP, closure of ATP-sensitive K^+^ (K_ATP_) channels, depolarization of the membrane which activates voltage gated calcium (Ca^2+^) channels, and elevated cytosolic free [Ca^2+^] activity which triggers insulin granule exocytosis and the release of insulin (1–3). Additional pathways (termed ‘amplifying pathways’) further enhance the release of insulin upon Ca^2+^ triggering. In type2 diabetes (T2D), these pathways are disrupted in the β-cell that can explain the insufficient release of insulin and altered glucose homeostasis (4). However, the exact mechanisms that underlie this dysfunction to the β-cells in diabetes are still unclear.

In islets, intra-cellular free Ca^2+^ activity ([Ca^2+^]) is tightly linked to electrical activity of the β-cell. Insulin is secreted in discrete pulses which is closely tied to oscillations in [Ca^2+^] and bursts of action potentials (5). Pulsatile insulin release allows for effective insulin action and glucose lowering (6–8). Connexin-36 (Cx36) gap junctions coordinate membrane potential between β-cells within the islet. As a result [Ca^2+^] oscillations are synchronized across the islet and insulin is released from the whole islet in discrete pulses (9). Thus, the regulation of [Ca^2+^] dynamics is a critical aspect to islet function.

While [Ca^2+^] is important for insulin granule release, its role as a secondary messenger can also affect gene transcription (10). For example, the serine/threonine phosphatase calcineurin (CaN) is activated by [Ca^2+^] which dephosphorylates its downstream target, nuclear factor of activated T-cell (NFAT). NFAT subsequently translocates to the nucleus to initiate transcription of a number of genes (11–14); thereby linking cell activity with gene transcription. This has critical importance in a number of systems to initiate remodeling, such as in activated T-cells or potentiating neuronal synapses (15). Within the islet NFAT regulates genes involved in β-cell function, including glucose sensing and granule formation (11, 13, 16, 17). NFAT also regulates genes involved in proliferation, with chemical activators of NFAT stimulating β-cell proliferation. While there is information regarding pathways in the β-cell activated by NFAT, it is unclear what physiological conditions within the β-cell lead to NFAT activation; particularly the link with stimulus-secretion coupling.

Here, we examine the link between the regulation of β-cell electrical activity and oscillatory [Ca^2+^], with NFAT activation and downstream transcription. We controlled the dynamics of islet electrical activity using either specific agonists/antagonists of ion channels together with optogenetics via light-activation of the cation channel ChR2. Using cell lines, mouse primary islets, and human primary islets, we determined the levels and dynamics of [Ca^2+^] under which NFAT would be activated and downstream gene transcription modified. We further tested whether altered Ca^2+^ regulation in models of T2D and human T2D impacted NFAT activation, and whether this can be normalized through exogenous control of [Ca^2+^]. Thus, overall we gain insight into the specific patterns of activity that regulate NFAT activation and gene transcription within the healthy and diabetic islet.

## Material and Methods

### Animals

All mouse experiments were performed in compliance with the University of Colorado Anschutz Medical Campus International Animal Care and Use Committee (IACUC). Mice were given water and food *ad libitum* and were housed in an environment with adequate temperature control with 12 h light/dark cycle. C57BL/6NHsd (C57BL/6) mice were obtained from Envigio. Rosa26-ChR2-YFP;Pdx-mice (18) were bred in house.

### Chemicals and Reagents

Diazoxide, Glibenclamide, Quercetin, bovine serum albumen (BSA), β-mercaptoethanol, tetraethylammonium (TEA), D-(+)-Glucose, and thapsigargin was purchased from Sigma-Aldrich (St. Louis, MO). FK506 was purchased from Enzo Life Sciences (Farmingdale, NY). SYBR Green qPCR Master Mix was purchased from ThermoFisher Scientific. Calcium indicator Rhod2-AM was purchased from AAT-Bioquest (Sunnyvale, CA). mCherry-NFATc3, GFP-NFATc1, GFP-NFATc2, GFP-NFATc3, GFP-NFATc4 constructs were a kind gift from Dr. Mark Dell’Aqua (University of Colorado Anschutz Medical Campus). ChR2-YFP plasmid was obtained from Addgene. Adenovirus particles containing mCherry-NFATc3 were prepared by the Diabetes, Obesity and Metabolism Institute, Icahn School of medicine at Mount Sinai. Cytosolic D3cpV FRET probe was a kind gift from Dr. Amy E. Palmer (University of Colorado Boulder).

### Cell Culture

Mouse insulinoma cells (MIN6) were grown in Dulbecco’s Modified Eagle’s Medium (DMEM, Corning) supplemented with 10% (v/v) fetal bovine serum (FVS) (Gibco), 100 U/mL penicillin, 100 μg/mL streptomycin (Gibco) and 60 μM freshly added β-mercaptoethanol. Cells were incubated at 37°C in 5% CO_2_ in a humidified controlled environment changing the media every 2-3 days. Once cells were approximately 90-95% confluent they were split and seeded onto 3.5cm glass bottom imaging dishes (35 mm petri-dishes, No1.5 cover glass, Corning, Ashland, MA) until they were approximately 75% confluent. At this point 2μg of mCherry-NFATc3, ChR2-YFP, and/or D3cpV plasmid DNA was transiently transfected using Lipofectamine 2000 (Invitrogen) mixed with OPTI-Mem (Gibco). This included incubating Lipofectamine 2000 and OPTI-Mem for 15min followed by the addition of DNA and incubating for 45 min. Media was changed prior to adding the transfection cocktail to MIN6 cells.

### Islet Isolation, Culture and Treatments

Islets were isolated from 12-16 week old mice, as previously described (19, 20). After pancreas dissection and digestion, and islet hand picking, islets were incubated for 3.5h at 37°C with 5% CO_2_ in a humidified environment in Roswell Park Media Institute (RPMI)-1640 media with 10% (v/v) FBS, 100U/mL penicillin, and 100μg/mL streptomycin. Islets were then transduced with 5μL of adenovirus titer 1.4*10^9^ plaque forming units (pfu) containing mCherry-NFATc3. After adenoviral incubation, islets were transferred to fresh RPMI-1640 media and incubated at 37°C overnight prior to start of experiments.

Islets from healthy and type 2 diabetic (T2D) human donors were obtained from the Integrated Islet Distribution Program (IIDP). IDs for human islets from donors without diabetes: AEFB055, AEGR353, AEHW247, AEHI491, AEIM290, AEJT193B, AEI1395, AEJQ100, and AEFO120; IDs for islets from donors with T2D: AEHL151, AEKH055, AELK219, AFBP453, AFBE414, and AFBK273. The average viability for healthy and T2D islets was 92% and 96% respectively. One set of islets from a donor with T2D was obtained from The University of Alberta Edmonton, Alberta, Canada (ID: R259). Upon arrival islets were incubated in CMRL 1066 medium (Corning) at 37°C, for 1h prior to mCherry-NFATc3 transduction. Transduction of mCherry-NFATc3 into healthy and T2D patient islets was done as stated above using mouse islets.

For pro-inflammatory cytokine treatment, mouse and healthy human islets were treated for 2h prior to start of experiment using vehicle control or a cytokine cocktail mixture specific for each mouse and human, consisting of 5ng/mL of recombinant interleukin-1β (IL-1β, R&D Systems, Minneapolis, MN), 10ng/mL recombinant tumor necrosis factor-α (TNF-α, R&D Systems), and 100ng/mL of recombinant interferon-γ (IFN-γ, R&D Systems) in RPMI-1640 media for 25min. Islets were imaged in solution supplemented with this cocktail. Mouse cytokine cocktail was used on MIN6 cells and incubated for 2h prior to the start of the experiment in DMEM media. MIN6, mouse and human islets were treated with 25 μM diazoxide or 1μM FK506 in DMEM or RPMI-1640 media for 2h prior to the start of the experiment and which was maintained constant in the imaging solution throughout the experiment. Human T2D islets were treated with 100 μM glibenclamide or 20 μM quercetin acutely post-start of experiment. MIN6 cells, healthy mouse and human islets as well as islets from donors of T2D were treated with 1μM thapsigargin, added to cells or islets 5 min after the start of the experiment.

### MIN6 and Mouse Islets Calcium Indicator Staining

To measure [Ca^2+^] changes in MIN6 cells we used the Förster Resonance Energy Transfer (FRET) D3cpV probe. MIN6 cells were seeded onto glass bottom imaging dishes and transfected with D3cpV, as described above. Imaging of MIN6 with D3cpV was done 24-48h post transfection under low and high glucose or incubated with either 25 μM diazoxide or 1 μM FK506 for 2 h prior to imaging and during imaging.

To measure [Ca^2+^] changes in MIN6 and β-cell specific ChR2-YFP expressing mouse islets we used the small molecule dye Rhod2-AM. MIN6 cells were seeded onto glass bottom imaging dishes and transfected with ChR2-YFP as described above. Prior to imaging, cells were incubated with 4 μM Rhod2-AM in imaging solution for 1-2 h at room temperature in a dark environment. Prior to imaging, cells were briefly washed with imaging solution and imaged in imaging solution supplemented with 2 mM glucose. Mouse islets containing ChR2-YFP β-cell were imaged two days post isolation. Islets were placed in a 35mm dish and stained with Rhod2-AM as described above using MIN6 cells. Islets were placed in a rotating table with slow to medium constant motion. After Rhod2-AM islet incubation islets were placed in glass bottom imaging dishes.

### Microscopy Imaging

Post-transfection/infection, cells or islets were imaged using BMHH imaging solution (125mM NaCl, 5.7mM KCl, 2.5mM CaCl_2_, 1.2mM MgCl_2_, 10mM HEPES) pH 7.4 supplemented with 0.1% BSA and 2mM glucose, 11mM glucose (for islets) or 20mM glucose plus 20mM tetraethylammonium (TEA) (for MIN6 cells, with TEA required to generate large coordinated Ca^2+^ oscillations (21–23). An Eclipse-Ti wide field microscope (Nikon) with a Nikon Plan Apo 20x/0.75 NA air objective or Nikon Apo LWD 40x/1.15 NA water immersion objective was used for islets and MIN6 cells, respectively. A Lambda 10-3 filter switcher and shutter controller was used with a Lambda LB-LS/30 Xenon arc lamp (Sutter Instruments). Images were acquired at 1 frame/15 s for 5 m at low glucose and 1 frame/15 s for 25 m after stimulation high glucose using a Andor CCD camera, with Nikon Elements software to operate the system. For optogenetic stimulation two pulse protocols were designed and imaging experiments were performed at low (2mM) glucose. For the first protocol, termed ‘slow pulse’ protocol, ChR2-YFP was activated at 1 frame/sec for 1 min followed 5 min of non-ChR2-YFP activation. This was repeated 5 times for the total duration of 30 min. For the second protocol, termed fast pulse protocol, ChR2-YFP was activated at 1 frame/sec for 30s followed 3 min of non-ChR2-YFP activation. This was repeated 8 times for the total duration of 30 min.

### RNA Extraction, qPCR and RNAseq

Islets isolated from 10-12 C57BL/6NHsd (C57BL/6) mice aged 8 weeks were used for RNA extraction. Islets were collected as stated above and given overnight recovery incubated in RPMI-1640 media. Islets were pooled together and batches of at least 250 islets were used per treatment. Batches were then treated with one of four treatments: 2mM glucose, 11mM glucose, 11mM glucose + diazoxide, or 11mM glucose + FK506 for 4 h. Batches treated with diazoxide or FK506 were respectively incubated with either 250μM diazoxide or 1μM FK506 at 2mM glucose for 1h prior to glucose elevation. Following treatment and incubation, islets were placed in a 1.5 mL tube, briefly spun down, and supernatant removed without disturbing islets. RNA was extracted using RNeasy® Micro Kit, and cDNA was synthesized using Omniscript RT Kit as per manufacturer’s instructions. For qPCR reaction 1 μL of cDNA was used per well and we used the following thermocycler conditions: Step 1 - 95°C for 10 m, Step 2 - 95°C for 15 s, and Step 3 - 60°C 45 s. Step 2 to step 3 were repeated a total of 60x. Forward and reverse primers used to perform qPCR for target genes of interest, as well as T_M_, can be found in **Table S1**. Other sets of islets were frozen for RNA extraction to be performed at a later date. Extracted RNA was snap frozen in liquid nitrogen and shipped to Novogene Corporation Inc (Sacramento, CA) for library preparation, RNA sequencing, and bioinformatics analysis. In each case for qPCR and RNAseq, RNA extraction was performed using RNeasy® Micro Kit, with Omniscript RT Kit used to obtain cDNA for qPCR (QIAGEN).

### Analysis

NFAT translocation in cells and islets was analyzed by drawing a region of interest in the cytosol and the nucleus of cells exhibiting fluorescence. GFP or mCherry fluorescent counts were collected frame by frame and the nuclear fluorescent signal was divided over the cytosolic fluorescent signal (^Nucleus^ ^fluorescence^/_Cytosolic fluorescence_ = Ratio). Time courses were normalize to the pre-stimulation signal ratio. For D3cpV experiments the FRET ratio (^FRET^/_CFP_ = FRET Ratio) was calculated by dividing FRET signal (YFP channel signal upon CFP excitation) by CFP signal (CFP channel signal upon CFP excitation), and plotted over time. Rhod2-AM and ChR2-YFP [Ca^2+^] experimental analysis in MIN6 cells was normalized to the pre-stimulation signal, followed by normalization to a double exponential regression curve (to account for photobleaching). Rhod2-AM and ChR2-YFP Ca^2+^ experimental analysis in mouse islets was also normalized to the pre-stimulation signal, followed by normalization to a linear regression curve.

Bioinformatics analysis of RNAseq was performed by Novogene Corporation Inc. This included a sample quality control to check for RNA Integrity Number (RIN), with only samples ≥6 further analyzed. After RIN test library preparation was performed, followed by sequencing by synthesis. Once sequence is obtained bioinformatics analysis was performed using FASTQ files to generate Fragments Per Kilobase of exon per Million (FPKM) readcounts. Heat maps were generated using FPKM readcounts, normalized to the FPKM at 2 mM glucose values, and expressed by the taking the *log_10_* (FPKM_norm_). Heat map limits were set to *−1* to *+1*.

qPCR was performed to obtain the C_t_-value from the target gene and the reference gene calculating the ΔC_t_. This was done by subtracting the target gene from the reference gene (HPRT C_t_ – Target Gene C_t_ = ΔC_t_). ΔC_t_ values were normalized to the ΔC_t_ values for 2 mM glucose treatment.

## RESULTS

### Glucose dependent electrical changes activates NFATc3 in MIN6 cells, mouse islets and human islets

To initially modulate electrical activity and measure the activation and translocation of NFATc3 we used the β-cell like mouse insulinoma MIN6 cell line (24). NFATc3 tagged with GFP was transiently transfected into MIN6 cells, and the ratio of nuclear to cytosolic fluorescence was monitored over time. At low (2mM) glucose GFP fluorescence was largely localized to the cytoplasm. Upon 20mM glucose (in combination with 20 mM TEA) there was a substantial increase in nuclear GFP fluorescence indicating NFATc3 activation and translocation to the nucleus (Figure 1A). Similar observations were made in mouse islets and human islets in which NFATc3-GFP was delivered via adenoviral infection: at low (2mM) glucose GFP fluorescence was largely localized to the cytoplasm whereas upon 11mM glucose there was a substantial increase in nuclear GFP fluorescence indicating NFATc3 activation (Figure 1B,C).

**Figure 1.**
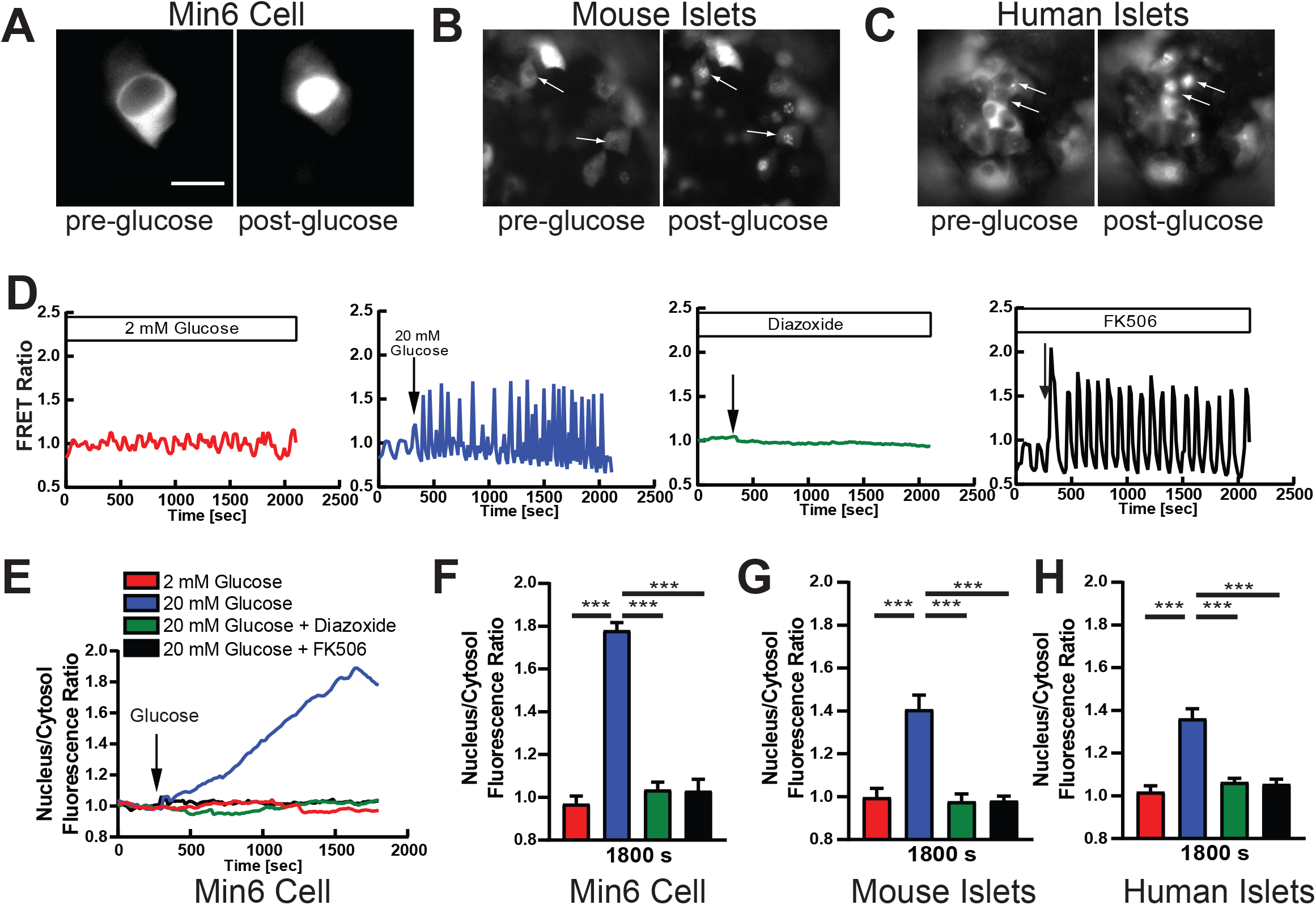
Glucose and membrane potential-dependent activation of NFATc3. **A)** Image of MIN6 cells transiently transfected with mCherry-NFATc3 at low glucose and 30 minute after addition of high glucose. **B)** as in A for mouse islets virally infected with mCherry-NFATc3. **C)** as in A for human islets from a healthy donor, virally infected with mCherry-NFATc3. **D)** Time-course of FRET ratio from D3cpV Ca^2+^ FRET sensor transfected into MIN6. Ca^2+^ changes were measured in four separate conditions: 2 mM glucose (red), 20 mM glucose (blue), 11 mM glucose + diazoxide (green), and 11 mM glucose + FK506 (black). **E)** NFATc3 translocation to the nucleus in MIN6 cells, for the conditions indicated in D, as measured by the ratio of nuclear to cytoplasmic fluorescence. Traces represent average of all experiments performed in MIN6 cells. **F)** mean nuclear-cytoplasmic mCherry-NFATc3 ratio 30minutes after treatment in MIN6 cells. **G)** as in F for primary mouse islets. **H)** as in F for primary human islets from a healthy donor. Statistical analysis was done using ANOVA with Tukey HSD post hoc test. *, **, and *** represent p=0.05, p=0.005, p=0.0005, respectively. Data in F averaged over n=5,11,8,4 plates respectively (18,31,21,8 cells); data in G averaged over n=3 mice (12,11,8,9 islets); data in H averaged over n=6 donors (n=8 for 2G, 11G+FK506; 28,21,24,33 islets)

Given elevated glucose stimulates membrane depolarization and [Ca^2+^] elevation, we next tested whether the NFATc3 activation that followed elevated glucose required membrane depolarization. Using a genetically encoded Ca^2+^ probe (D3cpV (25)), Ca^2+^ oscillations were absent at low glucose but as expected there are were robust increases in Ca^2+^ oscillations at elevated (20mM) glucose (**Figure 1D**). Upon treatment with the K_ATP_ activator diazoxide, which hyper-polarizes the cell, Ca^2+^ changes were abolished upon elevated glucose. In contrast upon treatment with CaN inhibitor FK506, the Ca^2+^oscillations at elevated (20mM) glucose were not perturbed. Using these treatment conditions, we tracked the activation of NFATc3, via its nuclear translocation, over time following glucose elevation. In MIN6 cells there was a progressive increase in the nuclear localization and activation of NFAT. This elevation was completely absent when maintained at low glucose (**Figure 1E**). Similarly, under both diazoxide treatment and FK506 treatment, glucose lacked any activation and nuclear translocation of NFATc3 (**Figure 1E**), indicating its activation is dependent on increases to the electrical activity. Following quantification significant nuclear translocation was only observed under elevated glucose, and not under low glucose, upon membrane hyperpolarization with diazoxide, or upon CaN inhibition with FK506 (**Figure 1F**). Similar observations were also made in primary mouse islets and human islets (**Figure 1G,H**) were nuclear localization and activation of NFAT was absent under low glucose, or elevated glucose under both diazoxide treatment and FK506 treatment.

NFAT consists of 4-isoforms, with each expressed in a tissue specific manner. All 4 NFAT isoforms are expressed in the β-cell (26, 27). To this end we tested if these isoforms were also activated via glucose and electrical activity. In MIN6 cells only NFATc2 and NFATc3 showed robust activation and nuclear translocation upon elevated (20mM) glucose (**Figure S1**). No significant nuclear translocation was observed with NFATc1 or NFATc4. Given the fact that in the β-cell high glucose activates both NFATc3 and Erk1/2 and both flank the insulin gene promoter to induce insulin transcription (28), we focused on NFATc3 activation.

The endoplasmic reticulum (ER) is a significant Ca^2+^ store in the cell (29, 30) and ER Ca^2+^ release can stimulate insulin secretion (31, 32). As such we tested whether ER Ca^2+^ release is sufficient to stimulate NFATc3 activation. In MIN6 cells, mouse islets and human islets at low glucose thapsigargin-induced ER Ca^2+^ release was sufficient to induce NFATc3 activation and nuclear translocation (**Figure S2**). Thus, elevated Ca^2+^ resulting from electrical activity, but also intracellular stores is sufficient to activate NFAT.

### Optogenetic-driven sustained electrical activity activates NFATc3

To further examine the dependence of NFAT activation and translocation on β-cell electrical activity we used optogenetics. ChR2 is a cationic channel that opens upon blue (450-480 nm wavelength) light, thus depolarizing the cell (33, 34). When expressed within the β-cell it allows tight control over the dynamics of electrical activity and [Ca^2+^] (35). Thus to stimulate islet electrical activity, independent of glucose, we illuminated primary mouse islets with β-cell specific expression of ChR2 (36) with different temporal patterns of light. We first measured the resultant [Ca^2+^] dynamics following either slow sustained pulses (1.5m ON/5m OFF) or more rapid pulses of light (30s ON/3m OFF), with an additional intermediate protocol (1m ON/5m OFF). In each case, following Rhod-2 staining, pulses of [Ca^2+^] were generated closely matching the pulse stimulation protocol (**Figure 2A,B** and **Figure S3A,B**). Furthermore, similar amplitudes were reached such that the time averaged [Ca^2+^] was similar across all three protocols (**Figure 2C** and **Figure S3C**). We next examined the resultant activation of NFAT through measuring the coincident translocation of mcherry-NFATc3 (**Figure 2D**), where use of mCherry avoids spectral cross-talk with the ChR2-YFP tag (37). Upon the slowest protocol there was a progressive elevation in NFAT activation and nuclear translocation following each sustained pulse of [Ca^2+^] elevation (**Figure 2D,E**). In contrast the more rapid pulse protocols did not elicit any NFAT activation and nuclear translocation (**Figure 2F** and **Figure S3D,E**). As such only the slow pulse protocol generated any significant NFAT activation and nuclear translocation compared to in the absence of ChR2 stimulation (**Figure 2G** and **Figure S3F**). Interestingly this slower protocol most closely matches physiological slow Ca^2+^ oscillatory dynamics within the islet under elevated glucose (5) This data demonstrates that the physiological pulsatile [Ca^2+^] dynamics are critical to activate NFATc3 in the β-cell.

**Figure 2.**
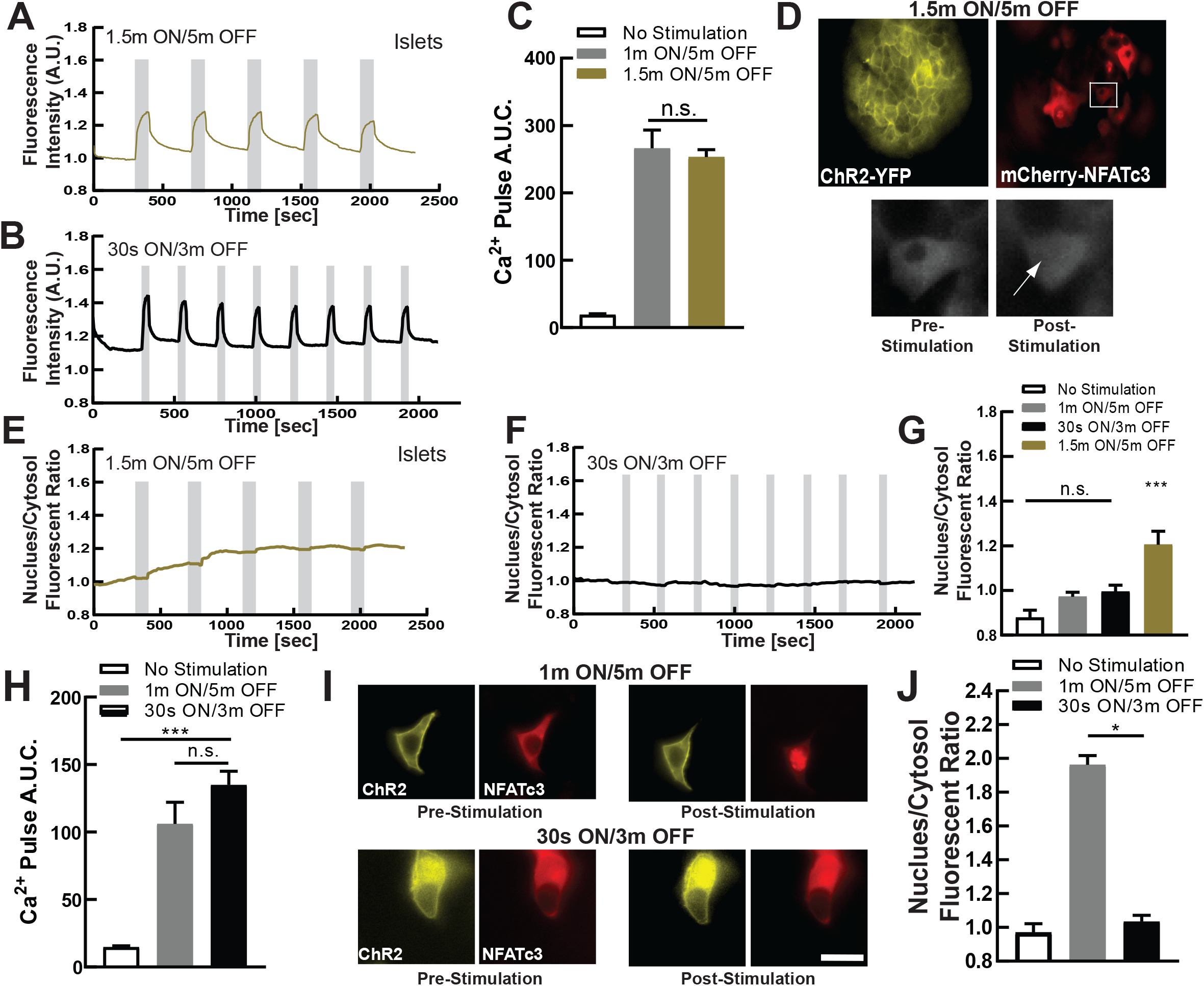
Dynamic Ca^2+^ changes via optogenetics activates NFATc3. **A)** mean Ca^2+^ changes in mouse islets expressing ChR2 in the beta-cell, at 2mM glucose following 1.5m ON/5m OFF optical stimulation protocol. Grey indicates duration of optical stimulation. **B)** as in A for 30s ON/3m OFF optical stimulation protocol. **C)** Area under the curve (AUC) for each stimulation pulse protocol, as well as time course lacking stimulation. **D)** representative images of ChR2-expresisng islet infected with mCherry-NFATc3, demonstrating NFATc3 translocation following optical stimulation (1.5m ON/5m OFF). **E)** Mean time-course of mCherry-NFATc3 nucleus/cytosol fluorescent ratio following the 1.5m ON/5m OFF optical stimulation pulse protocol. **F)** As in E for the 30s ON/3m OFF protocol. **G)** Mean nuclear-cytoplasmic mCherry-NFATc3 ratio 30minutes after optical stimulation. **H)** Area under the curve (AUC) of Ca^2+^ changes for stimulation pulse protocols (1m ON/5m OFF or the 30s ON/3m OFF), as well as time-course lacking stimulation, in MIN6 cells transfected with ChR2. **I)** representative images of ChR2-expresisng MIN6 cell transfected with mCherry-NFATc3, demonstrating NFATc3 translocation following optical stimulation. **J)** Mean nuclear-cytoplasmic mCherry-NFATc3 ratio 30minutes after optical stimulation in MIB6 cells. Statistical analysis was done sing ANOVA with Tukey HSD post hoc test. *, **, and *** represent p=0.05, p=0.005, p=0.0005, respectively. Data in C averaged over n=4 mice (n=6 for 1.5m/5m; 15,28 islets); data in G averaged over n=4 mice (n=6 for 1m/5m; 13,15 islets); data in H averaged over n=10,12 plates (29,27 cells); data in J averaged over n=10,24 plates (30,55 cells).

We further examined the link between [Ca^2+^] dynamics and NFATc3 activation in the β-cell using MIN6 cells transiently transfected with ChR2-YFP. Following Rhod2-AM staining similar [Ca^2+^] dynamics were elicited by each pulse protocol (**Figure S3G,H**), as in primary islets. For the intermediate (1m ON/5m OFF) and fast (30s ON/3m OFF) protocols similar amplitudes were reached. As such the time averaged [Ca^2+^] was similar across both protocols (**Figure 2H**). We were unable to utilize the slowest pulse protocol (1.5m ON/5m OFF) owing to the slower Ca^2+^ extrusion in MIN6 cells. In cells co-transfected with mCherry-NFATc3 we again examined the resultant activation and translocation of NFAT. During the slower intermediate protocol there was a progressive elevation in NFAT activation and nuclear translocation following each sustained pulse of [Ca^2+^] elevation (**Figure 2I** and **Figure S3I**). In contrast the more rapid fast pulse protocol did not elicit any NFAT activation and nuclear translocation (**Figure S3J**). As such only the slower intermediate pulse protocol generated any significant NFAT activation and nuclear translocation compared to in the absence of ChR2 stimulation (**Figure 2J**). Thus, a similar importance for pulsatile Ca^2+^ dynamics and NFATc3 activation were found in MIN6 cells, albeit with a shift in frequency dependence compared to primary islets.

### Altered electrical activity regulates RNA gene transcription of mouse islets

So far we have demonstrated that glucose stimulates NFAT activation and nuclear translocation, and this strictly requires elevated electrical activity. Furthermore, the physiological dynamics of electrical activity are critical to generate sustained [Ca^2+^] elevation required for NFAT activation and translocation. NFAT regulates many genes critical for β-cell function and proliferation (38, 39) and we next examined whether elevated electrical activity is important for NFAT-regulated gene expression. We incubated islets with either low or high glucose, as well as high glucose plus diazoxide to inhibit electrical activity or FK506 to inhibit CaN-NFAT signaling. Following 4 hours we isolated RNA to examine rapidly activated genes more likely to represent direct targets. We observed 57 genes were significantly upregulated by glucose and 55 genes significantly downregulated by glucose. Compared to these 112 glucose-regulated genes, 109 showed a significant reversal in gene expression upon diazoxide treatment. 40 of these genes showing overlap between the two conditions indicating their dependence on electrical activity. Of the 109 genes that showed a significant reversal in gene expression upon diazoxide treatment 19 showed a significant reversal in gene expression upon FK506 treatment, indicating their dependence on CaN-NFAT activity (**Figure 3A** and **Figure S4**). We next identified a subset of genes that have previously been associated with β-cell function and proliferation. These included immediate early genes such as the Fos family of genes, as well as Cdkl5, Dusp14, and Gpr161 genes (38) or Npas4 which is known to have cytoprotective properties in β-cells (40). Of these 18 genes that show the most significant increase in expression with elevated glucose and decrease in expression with glucose plus diazoxide, all but one also showed significant decrease in expression with FK506 (**Figure 3B**). Thus most ‘excitability-regulated’ genes that have a functional role in the β-cell are regulated by CaN-NFAT signaling. We performed analogous measurements in islets expressing ChR2 in the β-cell. Following 4 hours of pulsatile light stimulation (1.5m ON/5m OFF, **Figure 2A**) we again isolated RNA to examine genes activated rapidly by elevated electrical activity independent of glucose. We observed 25 and 31 genes that were significantly upregulated or downregulated, respectfully, following ChR2 stimulation. Of these genes, 56 ChR2-regulated genes, 22 were also observed to show glucose-regulated expression (**Figure 3C** and **Figure S4**). Similarly, of those genes with roles in the β-cell that most significantly changed upon glucose/diazoxide treatment, the vast majority also showed upregulation with ChR2 stimulation (**Figure 3D**). Thus, there are a large number of genes in which RNA levels are altered in response to changes to the electrical activity of the β-cell, dependent on CaN-NFAT signaling.

**Figure 3.**
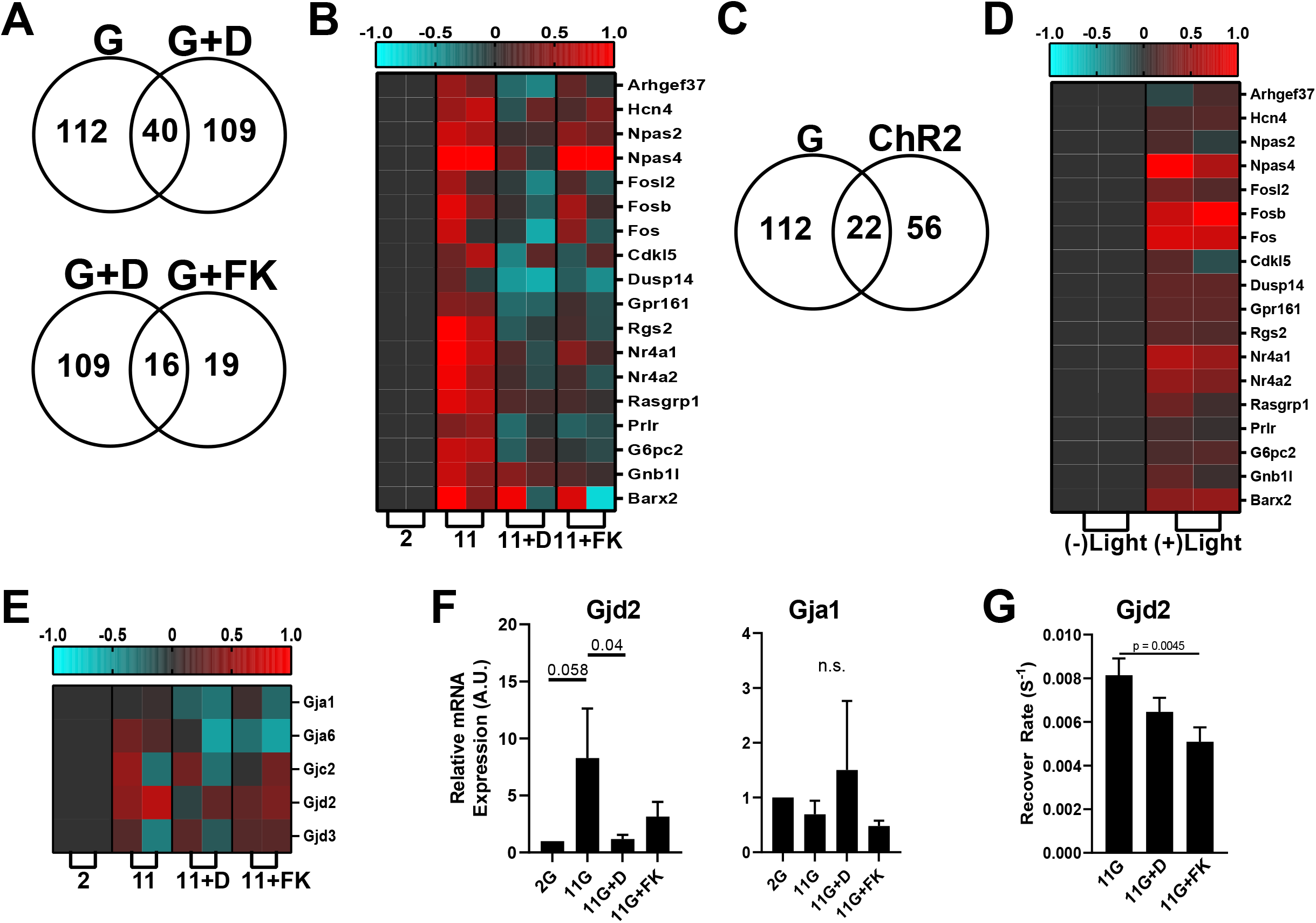
Acute gene transcription changes following elevated electrical activity. **A)** Venn diagram showing numbers of genes that are differentially expressed between glucose stimulation and application of K_ATP_ opener diazoxide upon glucose stimulation (upper) or between diazoxide application and fK506 application, upon glucose stimulation (lower). **B)** heat map depicting genes previously identified to regulate beta-cell proliferation and insulin secretion, following stimuli applied in A. Displayed is the subset of genes that shows the highest expression changes across the stimuli applied. **C)** Venn diagram showing numbers of genes that are differentially expressed between glucose stimulation and ChR2 optical stimulation. **D)** heat map depicting genes identified in B, following stimuli applied in C. **E)** heat map depicting connexin genes, following stimuli applied in A. **F)** qPCR of Gjd2 (Cx36) and Gja1 (Cx43) expression, **G)** fluorescence recovery after photobleaching measurements of gap junction permeability. Shown is data collected in islets treated with elevated glucose alone or upon diazoxide. Statistical analysis was done using ANOVA with Tukey HSD post hoc test. p values are indicated. ‘n.s.’ refers to p>0.2. RNAseq data in A-E represented from 2 independent experiments. Data in F averaged over n=5 experiments; data in G averaged over n=8 experiments.

*GJD2* encodes Connexin-36 (Cx36) which forms gap junction channels that electrically couple β-cells within the islets and coordinate Ca^2+^ regulation and insulin release across the islet (41). While not among the genes most highly upregulated by glucose, we observed that *GJD2* expression significantly increased upon glucose elevation, which was reversed by both diazoxide and FK506 (**Figure 3E**) indicating another gene in which expression is rapidly regulated by electrical activity via CaN-NFAT signaling. No other gap junction gene expressed within the islet showed such regulation. qPCR analysis supported these findings indicating glucose elevates *GJD2* expression (encoding Cx36) which is reversed by diazoxide and FK506, whereas *GJA1* (encoding Cx43), the other major connexin expressed within the islet, shows no such regulation (**Figure 3F**). Furthermore following 24h culture at elevated glucose, both diazoxide and FK506 significantly reduced functional gap junction permeability within the islet (42) (**Figure 3G**). Thus the elevation in *GJD2* expression following glucose-stimulated electrical activity and CaN-NFAT signaling is functionally relevant.

Previous work has established several genes involved in islet function and islet development/proliferation are regulated by CaN-NFAT signaling. RNAseq analysis (**Figure S5**) or qPCR analysis (**Figure S6**) each showed that these genes were not robustly increased acutely by elevated glucose nor decreased by diazoxide or FK506 treatment. An exception was that both FoxM1 and Hnf4alpha both showed significant changes in expression under elevated glucose which recovered upon diazoxide or FK506 treatment, indicating these genes may be acutely regulated by elevated electrical activity. Similarly, the NFAT subunits themselves also showed little acute changes in gene expression under these conditions (**Figure S4 and S5**). Thus while a number of genes important to β-cell function and development are acutely regulated by electrical activity and CaN-NFAT signaling, these are distinct from those identified in prior studies to be chronically regulated by CaN-NFAT signaling.

### Decline and restoration of NFATc3 activation in diabetic islets

Diabetic conditions disrupt [Ca^2+^] in the β-cell. This includes altered ER Ca^2+^ handling (43) but also gap junction communication, thus disrupting the dynamics of [Ca^2+^]. To test if NFATc3 is altered under diabetic conditions we first applied proinflammatory cytokines, which are elevated during diabetes, acutely to transduced mouse and human islets. Treatment with proinflammatory cytokines disrupts [Ca^2+^] elevation and dynamics (9, 44, 45). Following 2h incubation with proinflammatory cytokines, in primary mouse islets NFATc3 activation and translocation is blunted compared to untreated islets (**Figure 4A,B**). Similar blunting to NFATc3 activation and translocation was observed in primary human islets treated with proinflammatory cytokines (**Figure 4C**), and in MIN6 cells treated with proinflammatory cytokines (**Figure S7A,B).** Therefore, under acute diabetogenic condition NFAT activation is dysregulated in the β-cell.

**Figure 4.**
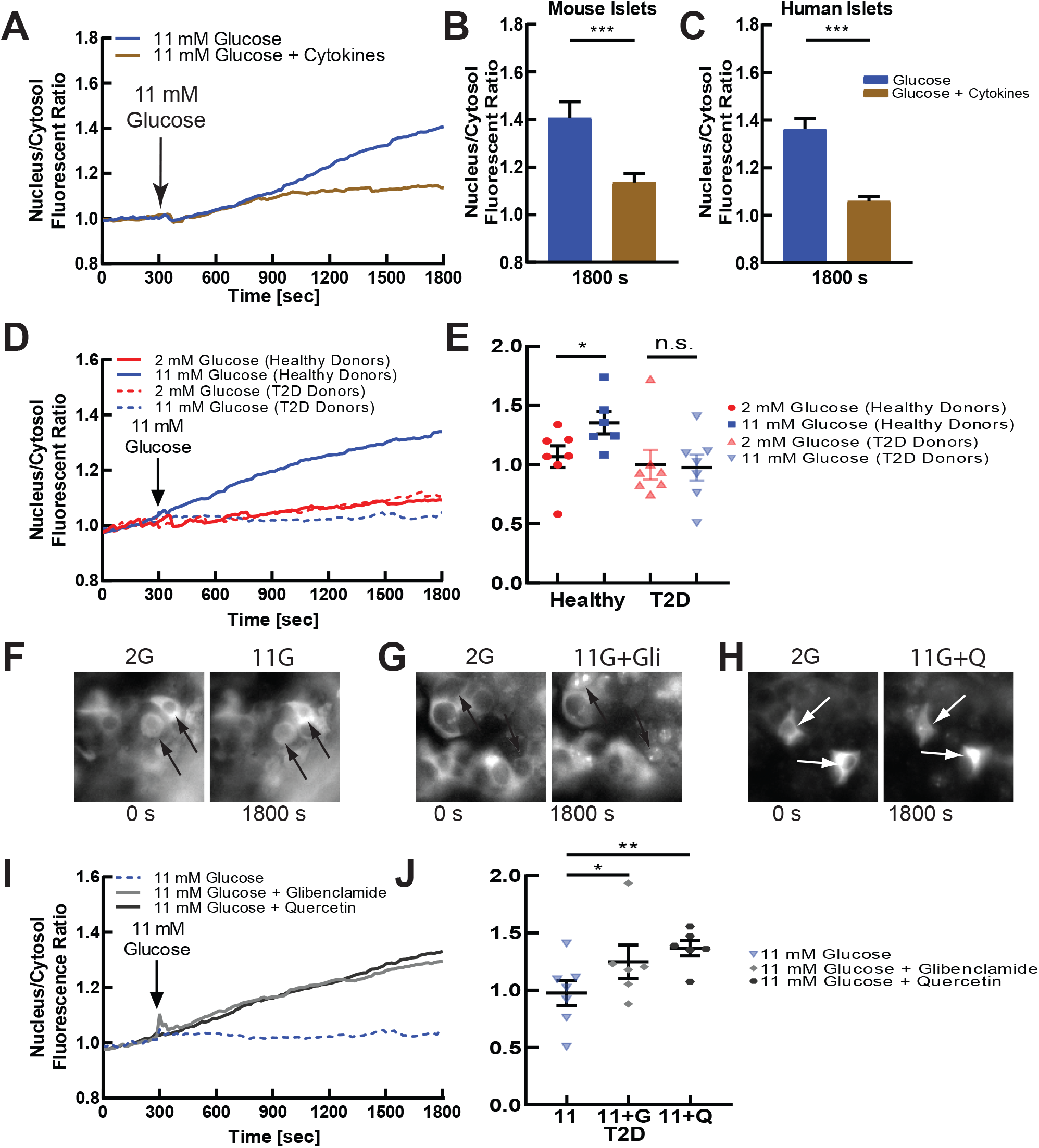
NFATc3 activation is diminished in diabetes due to diminished electrical activity. **A)** mean NFATc3 nuclear translocation over time, as measured by the ratio of nuclear to cytoplasmic fluorescence of mCherry-NFATc3, in untreated (blue) or cytokine-treated (brown) mouse islets, following elevated glucose. Cytokine treatment contains 0.005 μg/mL IL1-β, 0.01 μg/mL TNF-α, and 0.1 μg/mL IFN-γ for 2 h in DMEM or RPMI-1640 media. **B)** mean nuclear-cytoplasmic mCherry-NFATc3 ratio 30minutes after treatment in mouse islets, as in A. **C)** as in B for human islets from healthy donor treated with proinflammatory cytokines. **D)** mean NFATc3 nuclear translocation over time measured in human islets from healthy donors and donors with T2D, at 2mM glucose or following stimulation with 11mM glucose. **E)** nuclear-cytoplasmic mCherry-NFATc3 ratio 30minutes after treatment in human islets, as in A. Each dot represents the average all cells and islets from a specific healthy donor or donor with T2D. **F)** representative images of mCherry-NFATc3-infected human islet under low (2mM) and high (11mM) glucose. **G)** As in F under elevated (11mM) glucose or elevated glucose with Glibenclamide (Glib). **H)** As in F under elevated (11mM) glucose or elevated glucose with Quercetin (Q). **I)** mean NFATc3 nuclear translocation over time, under elevated glucose alone or including Glibenclamide (Glib) or Quercetin (Q). **J)** mean nuclear-cytoplasmic mCherry-NFATc3 ratio 30minutes after treatments in F-H. Each dot represents the average over all cells and islets from a specific donor with T2D, treated with the agents indicated. Statistical analysis was done using ANOVA with Tukey HSD post hoc test. *, **, and *** represent p=0.05, p=0.005, p=0.0005, respectively and n.s. means no significance. Data in B averaged over n=3 mice (12 islets); data in C averaged over n=8 donors (30 islets); data in E representative of n=6 healthy donors (21 islets) and n=7 donors with T2D (28 islets); data in J representative of n=6 donors with T2D (11,11,12 islets).

To test whether NFATc3 is dysregulated in more chronic cases of diabetes, we examined human islets obtained from either healthy donors or those with T2D. Human islets from donors with T2D showed diminished [Ca^2+^] under elevated glucose, but retained robust KCl-stimulated [Ca^2+^] (**Figure S8A**). Under low levels of glucose NFATc3 showed little activation and translocation in human islets from both healthy donors and T2D donors (**Figure 4D**). Under elevated glucose progressively elevated activation and translocation NFATc3 occurred in human islets from healthy donors (as already presented, **Figure 1**). However, in human islets from T2D donors little NFATc3 activation and translocation was observed (**Figure 4D**). Over a number of donors, significant NFAT activation was observed in healthy donor islets but not in T2D donor islets (**Figure 4E**). Specifically, human islets from 5/6 healthy donors showed NFAT activation and translocation under elevated glucose, whereas human islets from only 1/7 donors with T2D showed NFAT activation and translocation under elevated glucose. Thus, NFAT activation is significantly diminished in human T2D, correlation with the lack of robust glucose-stimulated [Ca^2+^] elevation.

To further test whether diminished NFAT activation and translocation in human T2D is a result of the diminished glucose-stimulated [Ca^2+^] elevation, we applied pharmacological stimulation of electrical activity and Ca^2+^ influx. Both the K_ATP_ channel inhibitor glibenclamide and voltage gated Ca^2+^ channel activator quercetin elevated [Ca^2+^] in human T2D islets (**Figure S8B,C**) indicating that the blunted glucose-stimulated [Ca^2+^] likely resulted from diminished glucose-sensing. We next tested whether these treatments would restore NFAT activation in T2D islets. Under elevated glucose, glibenclamide and quercetin led to progressively elevated activation and translocation of NFATc3 (**Figure 4F-I**). Over a number of donors significant NFAT activation was observed under both glibenclaide and quercetin, compared to elevated glucose alone, with quercetin eliciting a greater activation and translocation of NFATc (**Figure 4J**). These results show islets from T2D donors are electrically dysfunctional, likely as a result of altered glucose sensing and metabolism, thus diminishing NFAT activation. However pharmacological agents that restore electrical activity can restore activation of NFATc3.

## Discussion

Nuclear factor of activated T-cell (NFAT) is important in many tissues, playing a key role in linking cell activation to gene expression and cell remodeling. Within the islet NFAT regulates many genes involved in β-cell function, particularly in glucose sensing and insulin granule biogenesis (11, 16). NFAT is also a critical factor in β-cell proliferation, with inhibition of NFAT dephosphorylation and deactivation stimulating β-cell proliferation, including in human β-cells (46). While NFAT is activated under elevated glucose, little additional information is available for how NFAT is activated within the β-cell. Given the importance for electrical activity and Ca^2+^ influx in the β-cell, and the role for [Ca^2+^] elevation to activate NFAT in other systems, we sought to determine the link between the electrical activity of the β-cell and NFAT activation and downstream gene expression and how this is perturbed in diabetes when electrical activity is disrupted.

### Dynamic β-cell electrical activity regulates NFAT activation

We demonstrated that elevated electrical activity and [Ca^2+^] was required for NFAT activation: upon membrane hyper-polarization NFAT activation by glucose was completed abolished across all models examined. Furthermore, increased electrical activity was sufficient to activate NFAT, as demonstrated upon ChR2 stimulation at low glucose. While Ca^2+^ is long-known to activate NFAT, and elevated glucose in the β-cell is known to activate NFAT, this central role for β-cell electrical activity was previously not established. Importantly we observed this dependence on electrical activity only for c2 and c3 subunits, where the c2 subunit has also been linked to multiple aspects of β-cell function (11, 16).

Unexpectedly we also demonstrated that the dynamics of electrical activity and [Ca^2+^] also are important for NFAT activation: only slow sustained oscillations in [Ca^2+^] were sufficient for NFAT activation in both primary mouse islets and MIN6 cells. This was independent of Ca^2+^ ‘dose’ as similar time-averaged [Ca^2+^] was observed. This frequency-dependence is surprising as in other systems, such as chemical synapses, NFAT lacks frequency dependence (as opposed to CaMKII for example) (47). However, in chemical synapses the frequency varies with much faster time scales (seconds), whereas we see frequency dependence of bursts on a slower time scale (minutes). Furthermore, studies have shown [Ca^2+^] oscillations are more effective at inducing gene expression in T cells via NFAT compared to steady state levels (48). Nevertheless, the origin of this frequency dependence within the β-cell remains unclear. We speculate that it may originate from either some cooperativity that may only present on a minute time-scale, or that NFAT rephosphorylation and deactivation is negatively regulated by Ca^2+^-dependent processes. Determining this mechanisms will be important. Nevertheless, in revealing this unexpected frequency dependence, the use of optogenetics was critical, and this reveals a new avenue for optogenetic approaches.

The glucose-activation of NFAT was also diminished under diabetic conditions. Likely, this was a result of diminished [Ca^2+^], as pharmacological elevation of [Ca^2+^] via K_ATP_ inhibition or Ca_V_ stimulation recovered NFAT activation. We and others have measured diminished [Ca^2+^] in multiple models of diabetes, with this disruption originating from altered glucose-sensing and mitochondrial dysfunction (9, 41, 49). However, [Ca^2+^] oscillations are also disrupted in models of diabetes, showing shorter pulse duration (50). Thus, given elevated [Ca^2+^] does still exist in diabetic islets, another possibility is that the altered frequency and pulse duration is impacting NFAT activation in diabetes. Irrespective of the precise cause, the dysregulated [Ca^2+^] in diabetes is diminishing NFAT activation. As such factors that can promote robust elevated [Ca^2+^] will be important to recover NFAT activation and downstream transcriptional regualtion.

### NFAT links cell activity with gene expression

We observed a number of genes were upregulated or downregulated by conditions that increase electrical activity and [Ca^2+^] within the islet: both under glucose stimulation where up/downregulation was reversed by diazoxide-induced K_ATP_ opening; as well as following dynamic ChR2 stimulation. Many of the genes we identified were different from those previously identified to be NFAT regulated following either NFATc2 overexpression or CaN knockout (11, 16). The genes we identified included Ca^2+^ regulated genes that have been identified in other systems, such as Npas and Fos family (51–54). These genes are important to pathways underlying β-cell secretion, cytoprotection and proliferation (39, 40). The genes we identified also included those associated with cell cycle regulation and proliferation, consistent with the role for NFAT to regulate β-cell proliferation.

The difference between CaN-NFAT-regulated genes identified in this study, and those idnetiiexd in prior studies may result from the time-course of action. We examined genes rapidly upregulated by [Ca^2+^], whereas prior studies have utilized over-expression systems or gene knockouts, and thus reveal longer-term regulation. Thus, other transcription factors regulated by NFAT may be impacting these previously identified CaN-NFAT-regulated genes within the islet. Alternatively given that CaN is an effector for other molecules beyond NFAT, we may not be observing solely CaN-NFAT regulation. Nevertheless, we still do not see strong concordance between our gene expression results following pharmacological inhibition of FK506 and prior genetic knockout of CaN (16), suggesting the different time-scales being examined is the more likely explanation between the lack of correspondence between our study and prior results. We do observe some gene expression difference between those genes upregulated by elevated glucose which reversed upon hyper-polarization by diazoxide but lack upregulation by ChR2 stimulation. These differences may reflect an additional requirement of electrical activity, Ca^2+^ elevation and CaN-NFAT signaling, but also glucose acting via other pathways.

Indeed, when electrical activity and Ca^2+^ remain chronically suppressed via K_ATP_ mutation, islets show differing functional properties beyond electrical activity, even when the secondary effects of chronic hyperglycemia are factored out (55). Similarly if [Ca^2+^] remains chronically elevated, also via K_ATP_ mutation, numerous transcriptional changes occur (56). Pharmacologically-altered electrical activity impacts NFAT activation and downstream gene expression. Therefore the altered electrical activity and NFAT activation in diabetes will likely impact downstream, gene expression, and may underlie further islet dysfunction and diabetes progression. As such, restoring normal electrical activity may have broader impact beyond stimulating insulin release.

### Activity-dependent regulation of Cx36

Among those genes that we observed that are upregulated by elevated glucose and down regulated by membrane hyperpolarization induced by diazoxide was *GJD2*, which encodes Cx36 and forms gap junction channels within the islet. Thus, increased glucose enhances gap junction coupling, at least in part by electrical activity and Ca^2+^ influx. This is supported by functional data in which increased gap junction permeability was decreased by either membrane hyperpolarization or inhibition of CaN-NFAT signaling. Gap junction permeability and conductance is also enhanced following elevated glucose at much shorter time scales (e.g. <1h) than the time-scale on which we observed enhanced *GJD2* expression (>4h) (42, 57). Thus, while elevated glucose may stimulate increased *GJD2* transcription via elevated [Ca^2+^] and CaN-NFAT, glucose may also be required for post-translational regulation such as enhancing channel trafficking.

The electrical activity-dependent gap junction coupling that we observe may be important in the context of islet function: at elevated glucose, our results imply that increased electrical activity and [Ca^2+^] would lead to rapidly enhanced gap junction electrical coupling. In the context of cellular heterogeneity, this would ensure those more excitable cells with increased electrical activity and [Ca^2+^] would develop increased electrical coupling. A key role of electrical coupling is to coordinate different responses between cells (41, 58). As such we speculate that an increase in electrical coupling would increase the integration of the cell with the rest of the islet and thus overcome heterogeneous responses. For example, if a cell transitioned to become more proliferatively competent or entered the cell cycle its glucose responsiveness and insulin-secretory capacity would likely decline (59, 60). We speculate this may reduce electrical coupling in the less glucose-responsive cells, as has been observed for some cell sub-populations (59). This reduced electrical coupling would prevent the less-responsive cell negatively impacting the glucose responsiveness of the rest of the islet. Therefore, the activity-dependent regulation of Cx36 gap junction coupling within the islet may impact how functionally heterogeneous cells within the islet respond to dynamic changes in heterogeneity.

In summary, here we demonstrate that the electrical activity of the β-cell is critical to activation of NFAT, its nuclear translocation, and downstream gene transcription; particularly for a set of Ca^2+^ activated genes and other NFAT-regulated genes. This includes *GJD2* and the formation of Cx36 gap junction channels. The dysregulated electrical activity and [Ca^2+^] in diabetes leads to diminished or absent glucose-regulated NFAT activation, which can be rescued by pharmacological stimulation of electrical activity. This knowledge will be important to understand the regulation of NFAT and its target genes under normal and pathophysiological conditions.

## Acknowledgements

We would like to thank the Islet Isolation Core Facility at the Barbara Davis Center for Diabetes for allowing us to perform pancreatic mouse isolation and digestion to obtain islets. We would also like to thank Dr. Mark Dell’Aqua in the Department of Pharmacology at the University of Colorado Anschutz Medical Campus for giving us the fluorescent-protein-tagged NFATc3 constructs. RKPB receives support from NIH grants R01 DK102950, R01 DK106412 and JDRF grants 1-INO-2017-435-A-N. DGR receives support from NIH grant F31 DK121488.

## Data availability

All data presented in this study is available upon request.

## Electronic Supplemental Materials

**Supplementary Figure S1.**
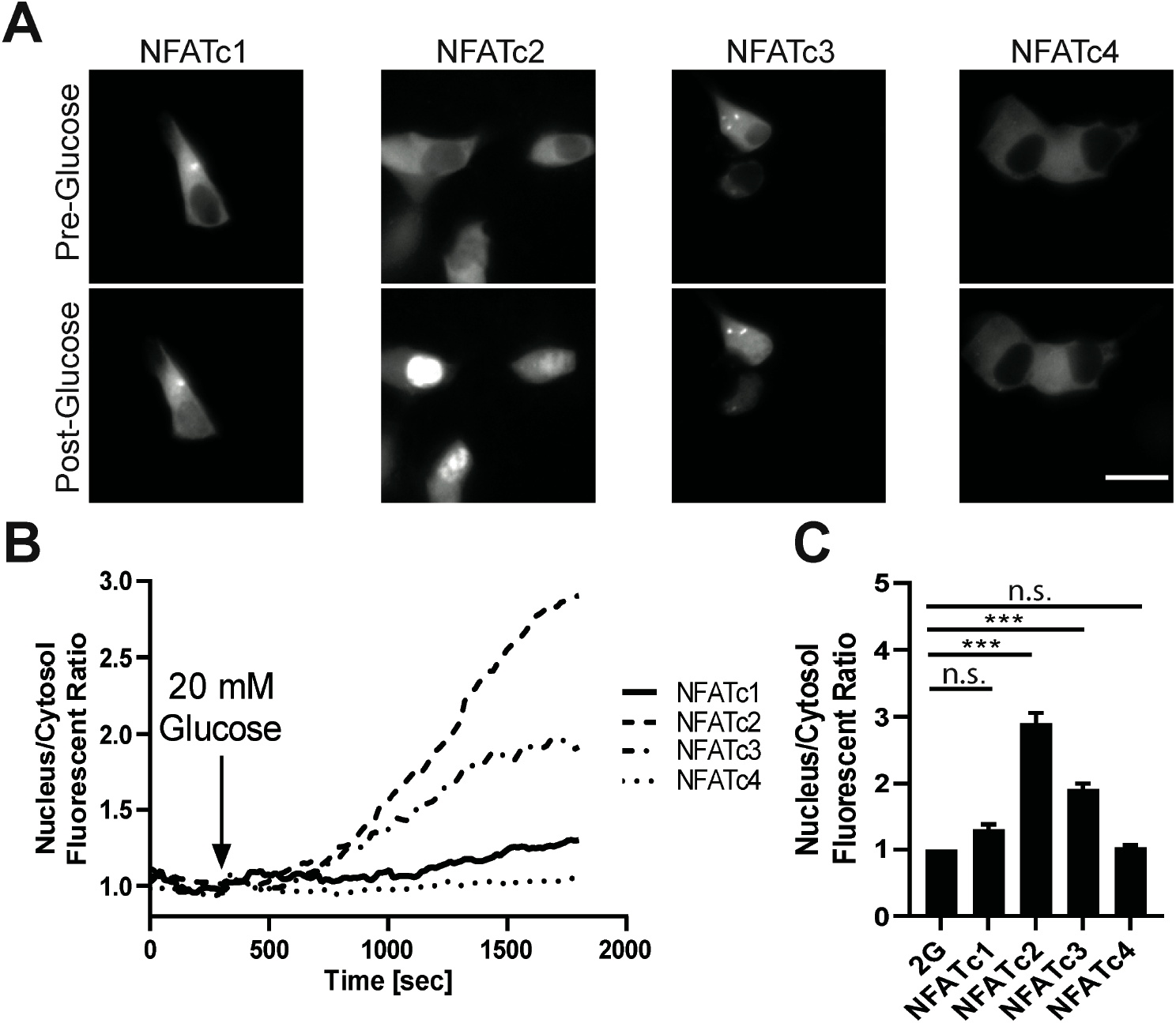
Glucose dependent activation of NFATc1-c4 in MIN6 cells. **A)** Images of MIN6 cells transiently transfected with mCherry-NFATc1,c2,c3,c4 at low glucose and 30 minute after addition of high glucose. **B)** Time-course for the ratio of nuclear to cytoplasmic fluorescence of NFATc1-4. Traces represent average of all experiments performed in MIN6 cells. **C)** mean nuclear-cytoplasmic mCherry-NFATc1-c4 ratio 30minutes after treatment in MIN6 cells. Statistical analysis was done using ANOVA with Tukey HSD post hoc test. *** represent p=0.0005, respectively and n.s. means no significance (p>0.2). Data in C averaged over n=4 plates (6,35,15,27 cells).

**Supplementary Figure S2.**
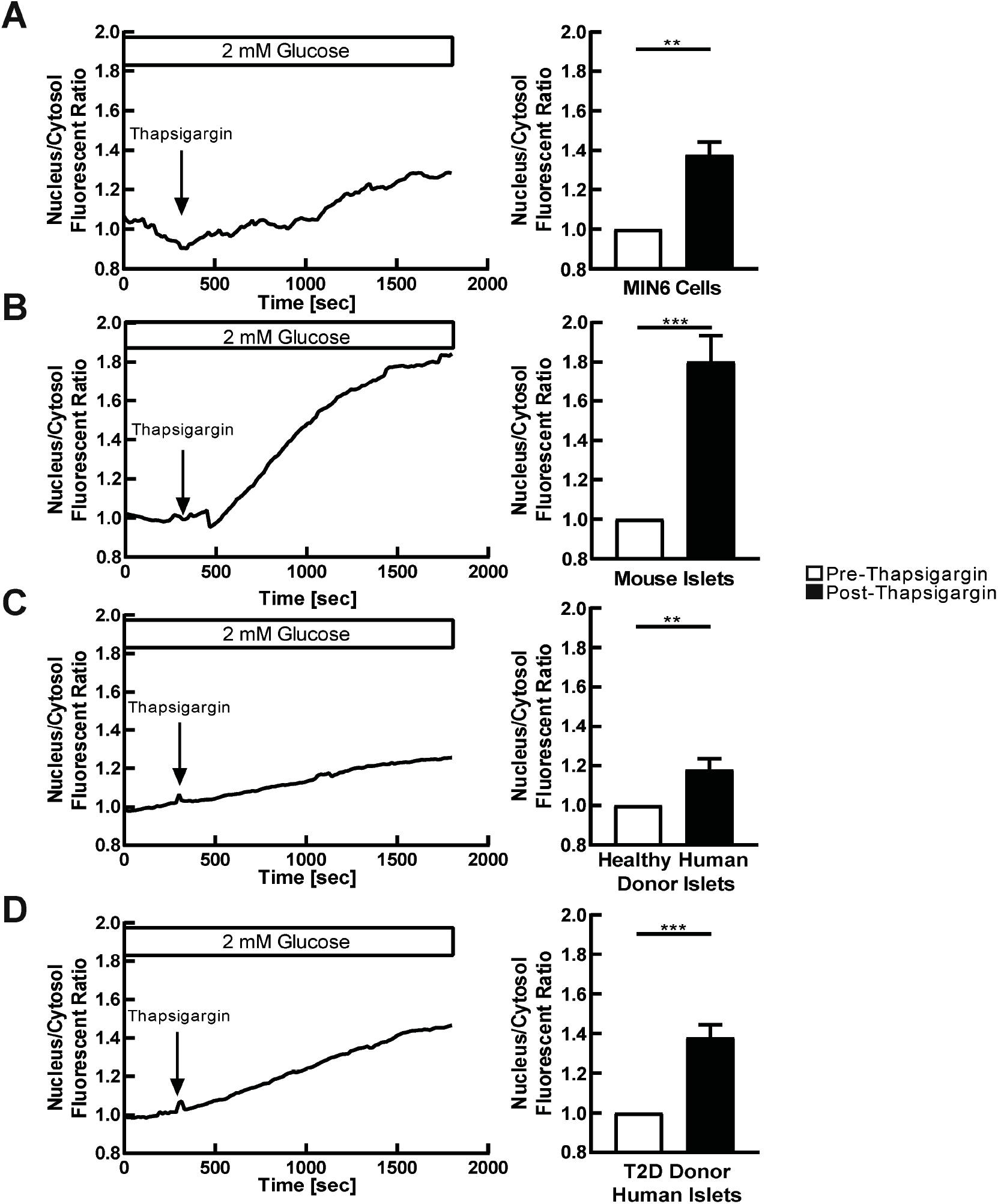
ER Ca2+ release activates NFATc3. **A)** Mean time-course for the ratio of nuclear to cytoplasmic fluorescence of MIN6 cells transiently transfected by mCherry-NFATc3, at 2 mM glucose and thapsigargin after addition of thapsigargin (left); together with mean nuclear-cytoplasmic mCherry-NFATc3 ratio before and 30min after thapsigargin addition (right). **B)** As in A for primary mouse islets infected with mCherry-NFATc3. **C)** As in A for human islets from healthy donors, infected with mCherry-NFATc3. **D)** As in A for human islets from donors with T2D, infected with mCherry-NFATc3. Statistical analysis was done using ANOVA with Tukey HSD post hoc test. *, **, and *** represent p=0.05, p=0.005, p=0.0005, respectively. Data in A-C averaged over n=8 plates (18 cells), n=2 mice (7 islets), n=8 healthy donors (31 islets) and n=8 donors with T2D (n=28 isletS).

**Supplementary Figure S3.**
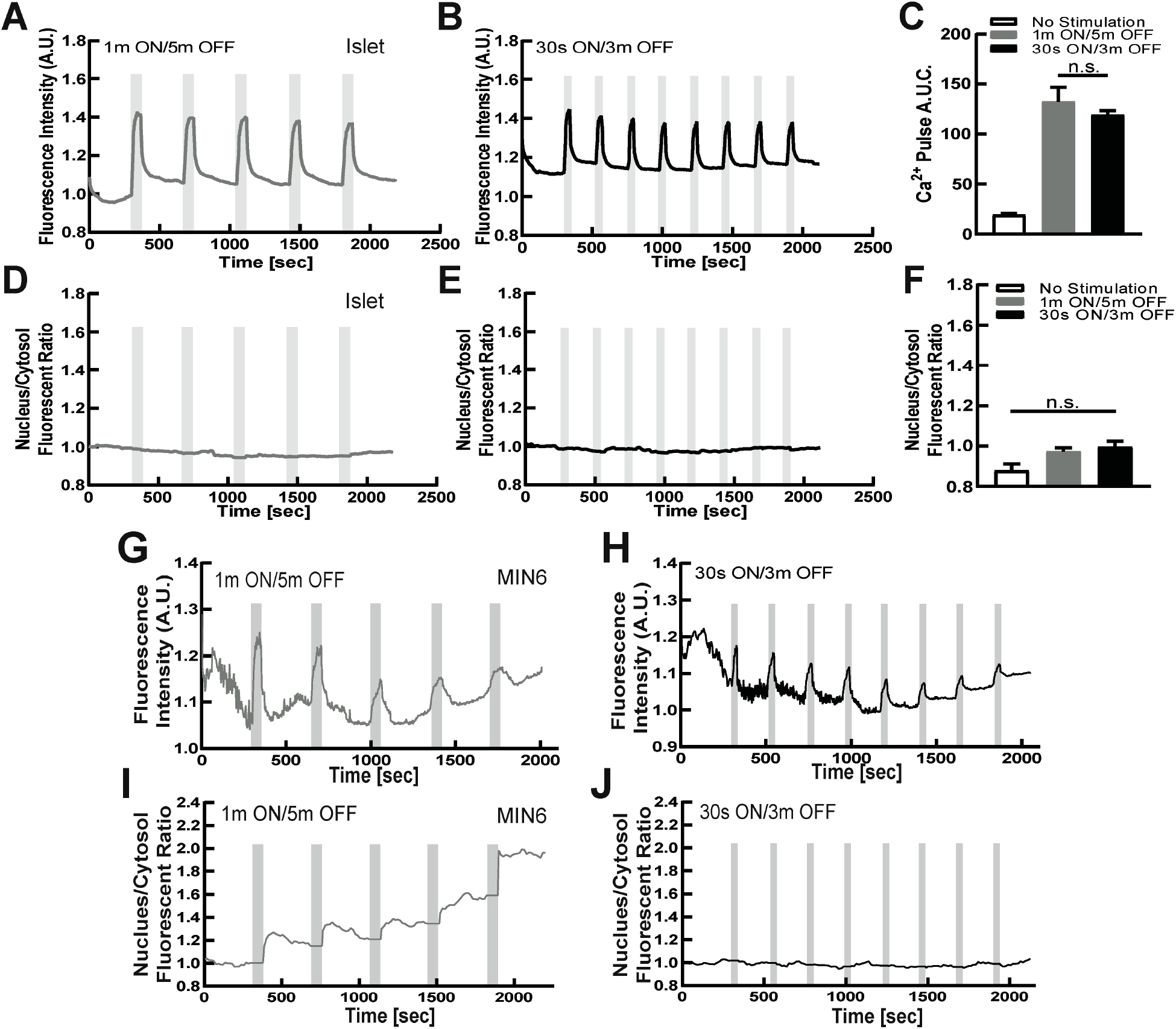
Dynamic Ca^2+^ changes via optogenetics activates NFATc3. **A)** mean Ca^2+^ changes in mouse islets expressing ChR2 in the beta-cell, at 2mM glucose following 1m ON/5m OFF optical stimulation protocol. Grey indicates duration of optical stimulation. **B)** as in A for 30s ON/3m OFF optical stimulation protocol. **C)** Area under the curve (AUC) for each stimulation pulse protocol, as well as time course lacking stimulation. **D)** Mean time-course of mCherry-NFATc3 nucleus/cytosol fluorescent ratio in ChR2-expressing islets following the 1m ON/5m OFF optical stimulation pulse protocol. **E)** As in D for the 30s ON/3m OFF protocol. **F)** Mean nuclear-cytoplasmic mCherry-NFATc3 ratio 30minutes after optical stimulation. **G)** mean Ca^2+^ changes in MIN6 cells transiently transfected with ChR2, at 2mM glucose following 1m ON/5m OFF optical stimulation protocol. Grey indicates duration of optical stimulation. **H)** as in G for 30s ON/3m OFF optical stimulation protocol. **I)** Mean time-course of mCherry-NFATc3 nucleus/cytosol fluorescent ratio in MIN6 cells, following the 1m ON/5m OFF optical stimulation pulse protocol. **J)** As in I for the 30s ON/3m OFF protocol. Statistical analysis was done using ANOVA with Tukey HSD post hoc test. Data in C,F= averaged over n=4 mice (15, 14 islets).

**Supplementary Figure S4.**
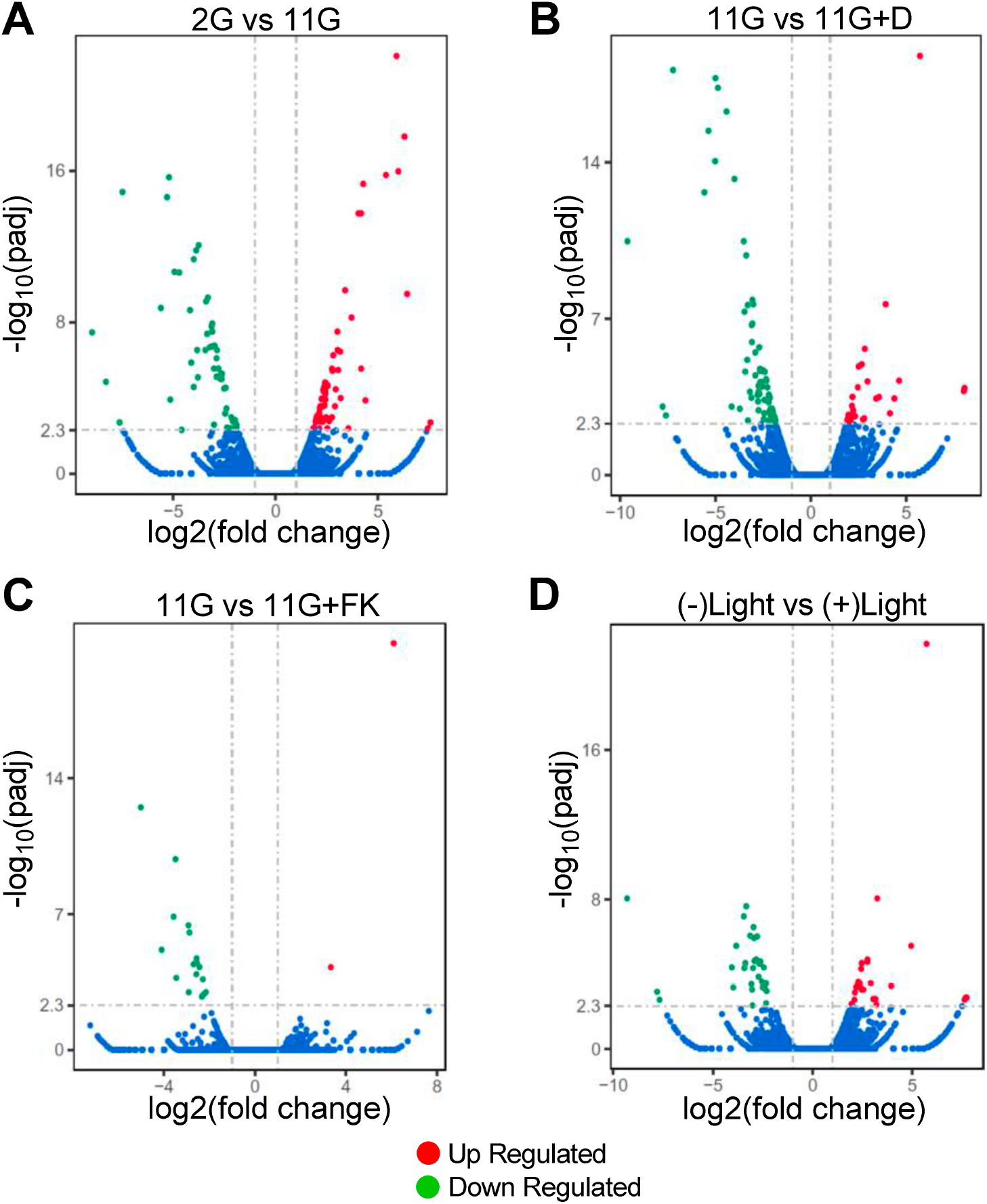
Changes in gene expression upon modulated electrical activity and Can-NFAT signaling. **A)** Volcano plot indicating changes in gene expression following RNAseq analysis of mouse islets treated with either 2 mM glucose (2G), or 11 mM glucose (11G) for 4h. Green indicates significant downregulation, red indicates significant upregulation. **B)** As in A for mouse islets treated with either 11 mM glucose (11G) or 11 mM glucose + diazoxide (11G+D) for 4h. **C)** As in A for mouse islets treated with either 11 mM glucose (11G) or 11 mM glucose + FK506 (11G+FK) for 4h. **D)** As in A for ChR2-expressing mouse islets at 2mM glucose treated with no light stimulation or pulsatile light stimulation for 4h.

**Supplementary Figure S5.**
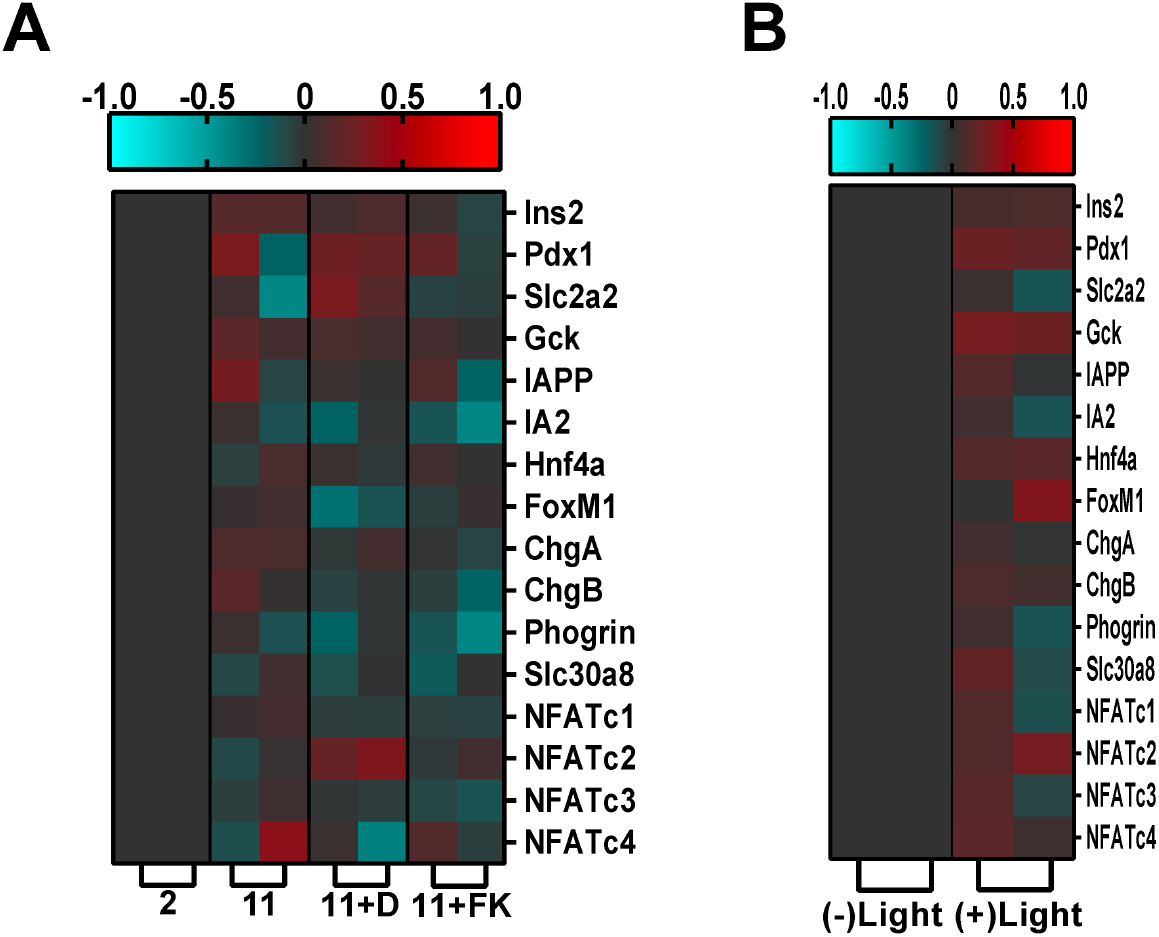
Acute gene transcription changes by RNAseq following elevated electrical activity. **A)**. Heat map depicting genes previously identified to regulate beta-cell proliferation and insulin secretion, following glucose stimulation, application of K_ATP_ opener diazoxide upon glucose stimulation upon application of CaN inhibitor FK506 upon glucose stimulation. B) As in A for CR2-expressing islets in the absence and presence of stimuli applied in A. Displayed is the subset of genes that shows the highest expression changes across an optical stimulation protocol.

**Supplementary Figure S6.**
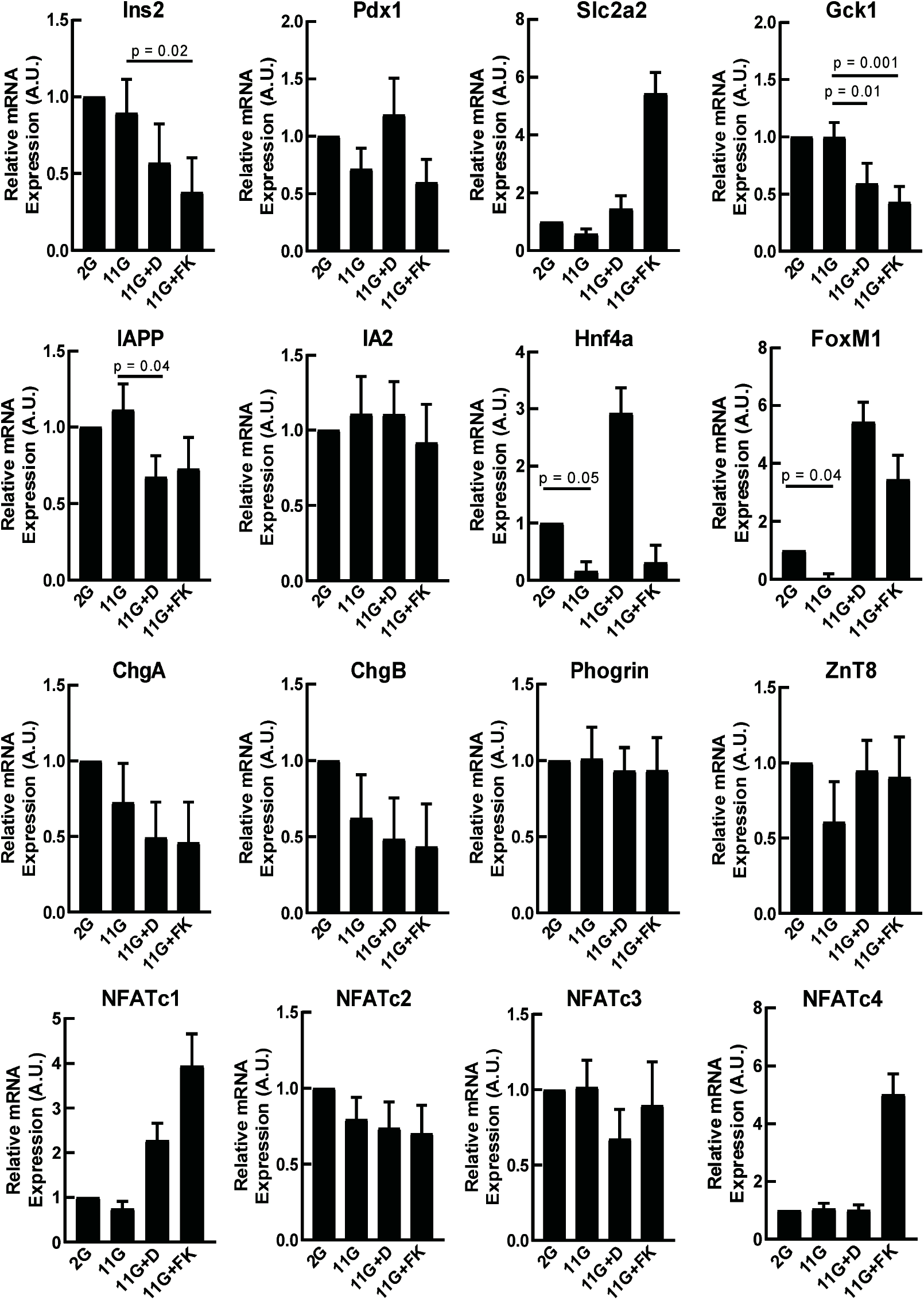
Acute gene transcription changes by qPCR following elevated electrical activity. qPCR measurements of gene expression for those genes genes previously identified to regulate beta-cell proliferation and insulin secretion indicated in Supplementary figure 5. Genbe expression follos glucose stimulation, application of K_ATP_ opener diazoxide upon glucose stimulation, or upon application of CaN inhibitor FK506 upon glucose stimulation. Statistical analysis was done using ANOVA with Tukey HSD post hoc test. Data in F averaged over n=5 experiments.

**Supplementary Figure S7.**
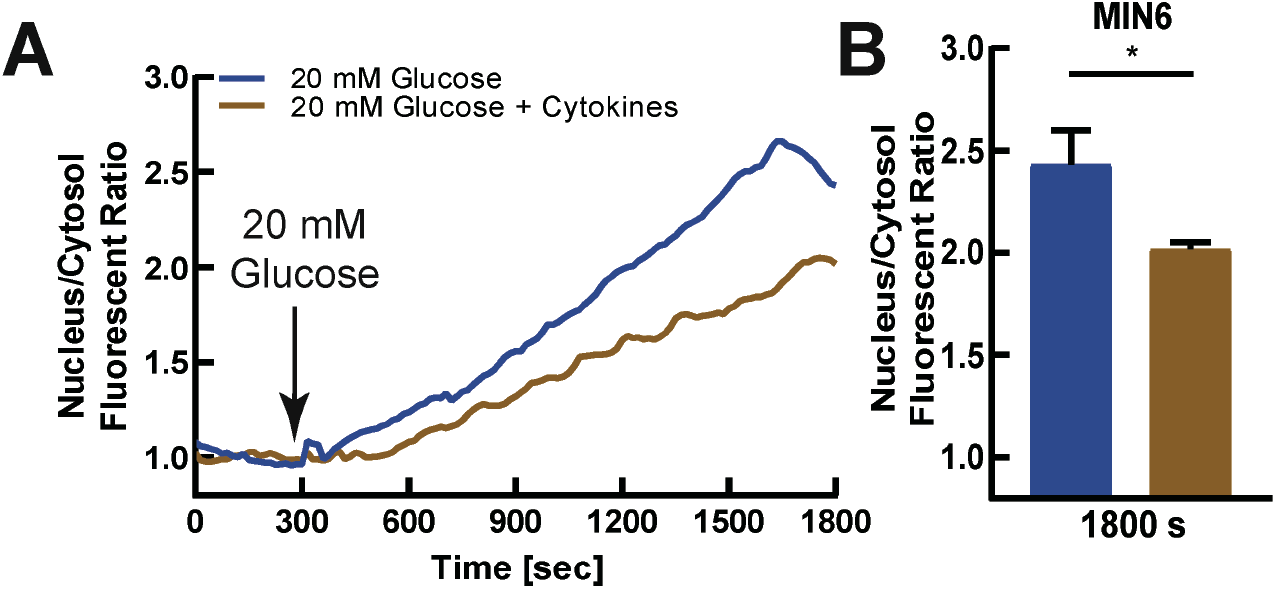
Diminished NFATc3 activation in MIN6 cells treated with proinflammatory cytokes. Mean NFATc3 nuclear translocation over time, as measured by the ratio of nuclear to cytoplasmic fluorescence of mCherry-NFATc3, in untreated (blue) or cytokine-treated (brown) MIN6 cells, following elevated glucose. Cytokine treatment contains 0.005 μg/mL IL1-β, 0.01 μg/mL TNF-α, and 0.1 μg/mL IFN-γ for 2 h in DMEM or RPMI-1640 media. **B)** mean nuclear-cytoplasmic mCherry-NFATc3 ratio 30minutes after treatment in MIN6 cells, as in A. Statistical analysis was done using ANOVA with Tukey HSD post hoc test. * represent p=0.05, Data in B averaged over n=5 plates (17 cells).

**Supplementary Figure S8.**
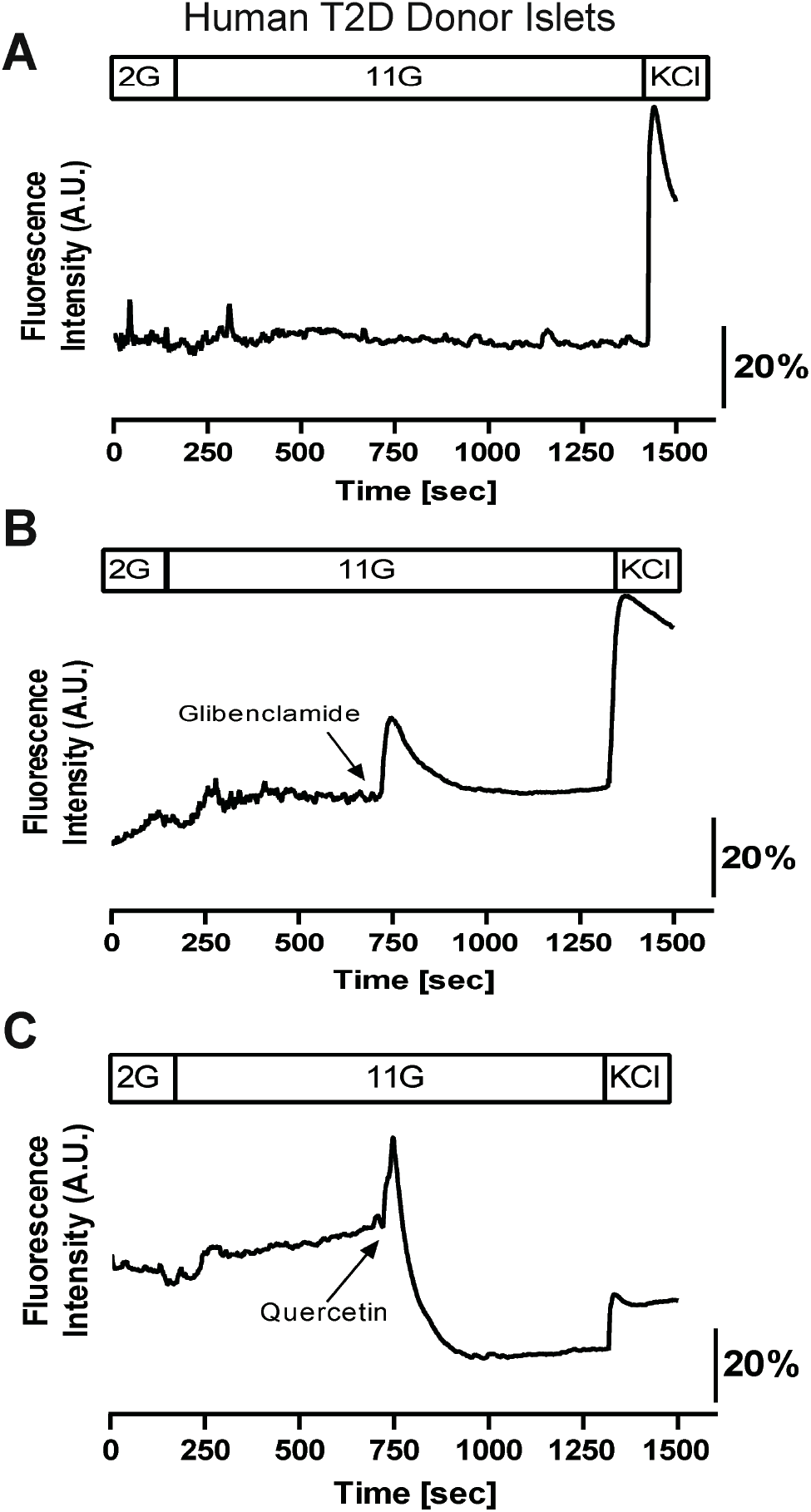
[Ca^2+^] response in islets from donors with T2D. **A)** [Ca^2+^] elevations, as indicated by Fluo4 fluorescence, in human islets obtained from donors with T2D. Treatments include sequential stimulation for 2 minutes at 2 mM glucose, 20 minutes at 11 mM glucose, and 3 minutes with 20 mM KCl. **B)** As in A, but 10 minutes post adding 11 mM glucose, glibenclamide was added acutely. **C)** as in A but 10 minutes post adding 11 mM glucose, quercetin was added acutely. Data representative of islets 3 donors with T2D (11 islets).

**Supplementary Table S1.**
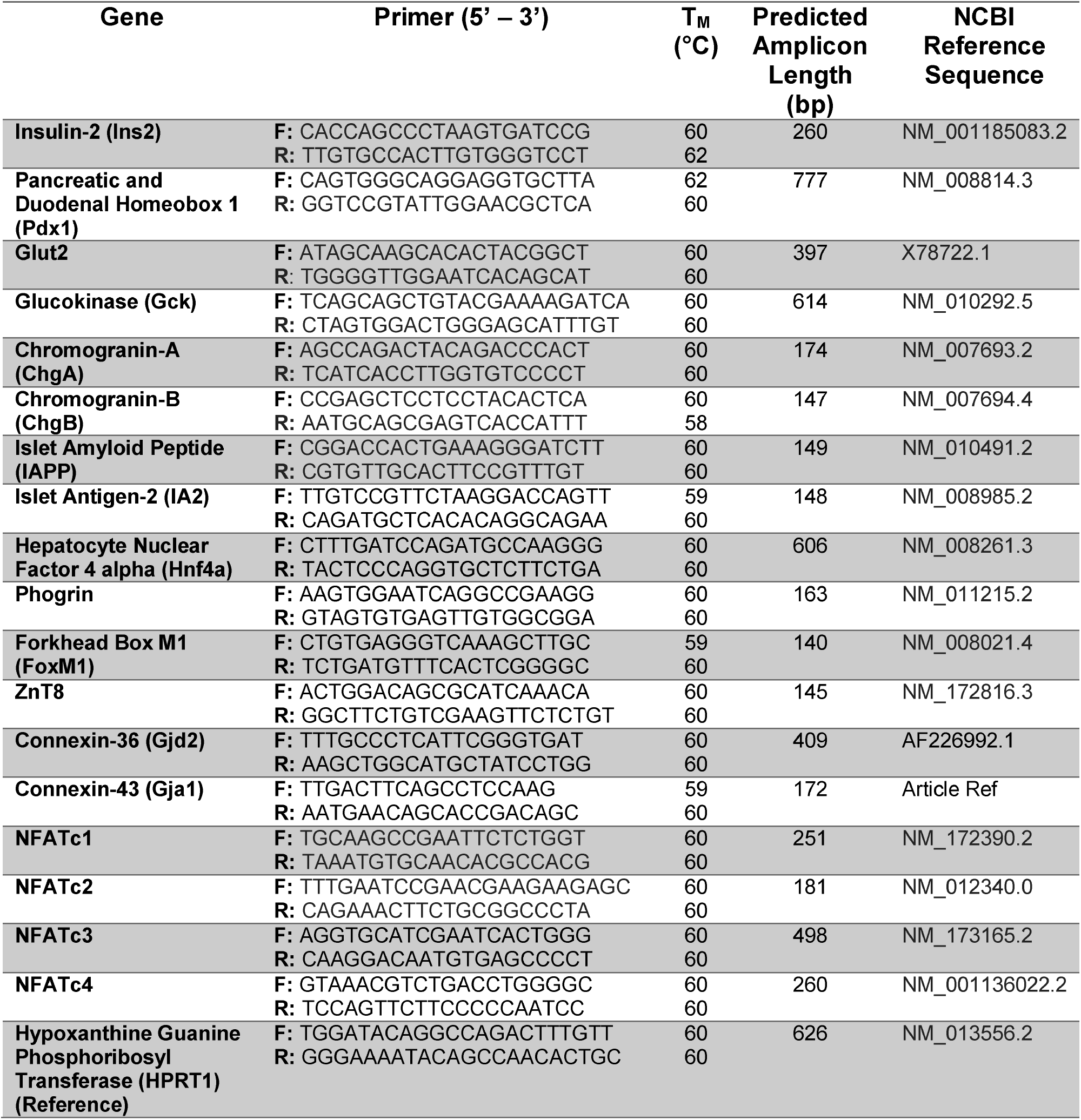
qPCR primers. Primers used to test specific genes involved in cell function, insulin granule formation, and NFATc1-NFATc4. Table shows primer name, sequence from 5’-3’, melting temperature, and predicted amplicon.

